# A DEEP LEARNING APPROACH TO ESTIMATING INITIAL CONDITIONS OF BRAIN NETWORK MODELS IN REFERENCE TO MEASURED FMRI DATA

**DOI:** 10.1101/2021.07.07.451431

**Authors:** Amrit Kashyap, Sergey Plis, Michael Schirner, Petra Ritter, Shella Keilholz

## Abstract

Brain Network Models (BNMs) are a family of dynamical systems that simulate whole brain activity using neural mass models to represent local activity in different brain regions that influence each other via a global structural network. Research has been interested in using these network models to explain measured whole brain activity measured via resting state functional magnetic resonance imaging (rs-fMRI). Properties computed over longer periods of simulated and measured data such as average functional connectivity (FC), have shown to be comparable with similar properties estimated from measured rs-fMRI data. While this shows that these network models have similar properties over the dynamical landscape, it is unclear how well simulated trajectories compare with empirical trajectories on a timepoint-by-timepoint basis. Previous studies have shown that BNMs are able to produce relevant features at shorter timescales, but analysis of short-term trajectories or transient dynamics as defined by synchronized predictions from BNM made at the same timescale as the collected data has not yet been conducted. Relevant neural processes exist in the time frame of measurements and are often used in task fMRI studies to understand neural responses to behavioral cues. Therefore, it is important to investigate how much of these dynamics are captured by our current brain simulations. To test the nature of BNMs short term trajectories against observed data, we utilize a deep learning technique known as Neural ODE that based on an observed sequence of fMRI measurements, estimates the initial conditions such that the BNM’s simulation is synchronized to produce the closest trajectory relative to the observed data. We test to see if the parameterization of a specific well studied BNM, the Firing Rate Model, calculated by maximizing its accuracy in reproducing observed short term trajectories matches with the parameterized model that produces the best average long-term metrics. Our results show that such an agreement between parameterization using long and short simulation analysis exists if also considering other factors such as the sensitivity in accuracy with relative to changes in structural connectivity. Therefore, we conclude that there is evidence that by solving for initial conditions, BNMs can be simulated in a meaningful way when comparing against measured data trajectories, although future studies are necessary to establish how BNM activity relate to behavioral variables or to faster neural processes during this time period.

## 2. Introduction

Brain Network Models (BNMs) represent whole brain activity as the coordination of many distinct neural populations that are connected via a structural network consisting of long-distance white matter tracts (Sanz Leon et al., 2015, Breakspear et al., 2017). Simulations of these network models are being compared to experimental measurements such as functional magnetic resonance imaging (fMRI). At this spatiotemporal scale in fMRI, the measured activity is thought to be an averaged property of the neural populations and occurs relatively slowly (<1Hz) compared to the faster neural information processes and the measured fMRI signal is thought to represent the coordination between brain regions over the structural network (Deco et al., 2009). BNMs have been able to reproduce properties observed in fMRI, especially during rest where the brain is not exposed to a structured experimental task or stimulus and the whole brain activity is thought to mostly arise from intrinsic network loops between cortical regions (Honey et al 2007, Cabral et al., 2011, Sanz Leon et al., 2015). Thus, BNMs have been used as a generative framework in order to analyze how local neural activity could translate to global coordination and how changes due to neural pathologies translate to observed aberrant dynamics (Sanz Leon et al., 2015, Ritter et al., 2013, Schirner et al., 2018, Saenger et al., 2017).

Current research does not utilize a single type of population model to construct BNMs. Rather, depending on the application and the underlying assumptions, a choice of neural mass models with different properties are selected and used to construct BNMs (Sanz Leon et al., 2015). For the most widely used application - replicating rs-fMRI - several models have been shown to reproduce time-averaged properties such as functional connectivity (FC), computed via cross correlation of long time-courses of pairs of brain regions (Cabral et al., 2017). These comparisons over long simulation windows to long rs-fMRI timeseries are made because the initial conditions are not well-defined in resting state as there is no set stimulus onset to use as a reference when comparing with the measured signal (Cabral et al., 2011). This has led to using different dynamic metrics computed over long time-windows on both simulated and measured data and then comparing the outputs of the metrics across modalities, rather than comparing the measured timeseries with the output of the models directly (Cabral et al., 2017, Kashyap & Keilholz 2019). BNM predictions at a time point by time point basis, is of interest to researchers as many neural processes observed in fMRI occur over these timescales, such as responses to task stimuli or aberrant responses due to neural pathologies. While faster processes in BNM’s have been examined in previous studies and how they compare with multimodal recordings such as EEG data (Schirner et al., 2018), but no study has investigated how well short-term trajectories, defined by a series of consecutive fMRI measurements, are being reproduced by our current BNMs. For such an experiment, we address the gap of solving for initial conditions relative to an observed trajectory for a given BNM, and then compare the synchronized predictions of the simulation with the observed timeseries.

We hypothesize that the BNMs that are better approximation of the underlying dynamical system in whole brain dynamics determined using traditional long-term measures, will evolve more closely to the measured rs-fMRI trajectories. An agreement of parameterization between the long-term metrics and differences in these short trajectories would support the evidence that BNM’s are simulating meaningful trajectories that can be compared directly with fMRI timeseries and can be later used to investigate neural processes during this timeframe.

The initial conditions are estimated, by utilizing a novel method developed in the Machine Learning community that utilizes a sequence of observations and a given dynamical system to output the initial conditions of the dynamical system that would be the closest fit to the current observed data trajectory. The technique, known as Neural Ordinary Differential Equations (ODE), uses a recurrent neural network (RNN) that keeps track of information from previous timepoints, in order to predict the initial conditions of a given dynamical system based on previous observations (Chen et al., 2019). The neural network model is trained via one step prediction, where from the estimated initial conditions and the known dynamical system predicts the next timestep via integration and compared with the true next step. The algorithm, therefore, regardless of the dynamical system, gives similar predictions over the first-time interval but the trajectories diverge over longer periods of integration due to differences in the dynamical system and become less dependent on initial RNN predictions.

A potential issue to this approach is that both the signal simulated as well as the measured rs-fMRI signal are thought to be produced by stochastic processes. This might affect the approaches’ ability to discriminate correctly between different BNM on their ability to simulate rs-fMRI. However, previous studies have shown that dynamic metrics are better than metrics computed over a long period of time to parameterize BNM, thus suggesting that even in shorter windows allows for discrimination between models (Kashyap & Keilholz 2019).

To test if this approach can correctly identify components of a known BNM, we used the Firing Rate Model (FRM) from Cabral et al., 2012 with the same equations and network regions as a candidate dynamical system to fit to the rs-fMRI data. The FRM is a linear model that defines the change of dynamics in a single neural population as a weighted sum of its network neighbors and applies an exponential decay term to prevent runaway excitation (Cabral et al., 2012). The model contains three components (global coupling, noise amplitude, structural matrix), which are varied independently, and a specific Neural ODE is trained for each variation to solve for the initial conditions. The results show that without noise, maximizing on accuracy over the short time window yields trivial BNMs that don’t depend on the structural connectivity. However, in the presence of noise the trend reverses and models with strong structural connectivity perform better than the models with weak or no network influence. Since the value of noise is not known, an additional parameter namely the structural connectivity was varied by slowly adding noise to the original connectivity, and the sensitivity due to this change in network was measured. The FRM that showed the greatest changes in accuracy due to perturbations of the structural connectivity had a parameterization that was in agreement to the one established using long term metrics.

In short, the manuscript shows that Neural ODE approach can be used to simulate FRM trajectories that can be directly compared with measured rs-fMRI data in a meaningful manner. The differences between parameterizations of the model with respect to rs-fMRI data observed during this timeframe are similar to those observed over longer simulations. Therefore, this tool can be used in the future to analyze BNM on shorter timescales to measured data such as for task fMRI. Moreover, it can be used as an unbiased metric to compare the signal directly with the models and aid our development of discovering more powerful models that recapitulate whole brain activity.

## 3. Methods

### 3.1 Overview

This section is organized by first describing functions that are used to fit to the rs-fMRI data, mainly BNMs but also certain null models that are used for comparison. The subsequent sections then describe how the Neural ODE algorithm is used to infer the initial conditions of a given dynamical system based on previous measurements. We describe our own implementation of the Neural ODE algorithm that was specifically designed to train on large amounts of imaging data (Chen et, al. 2018, GIT REF https://github.com/sagastra/fMRINeuralODE). The algorithm was validated using synthetic data from a simple spiral dynamical system described in detail in the supplementary sections 1-3. The subsequent sections after describing the algorithm, deal with the processing of experimental fMRI and DTI data used to construct the models. The final section outlines how the simulated trajectories are compared with empirical trajectories.

### 3.2 Brain Network Models

Brain Network Models are used as models for whole brain network activity. They combine a mathematical description of the intrinsic activity of a neural population with the global brain structure that coordinates the activity between populations. In order to construct a BNM, the structural network is first defined, usually based on a parcellation scheme that outlines which cortical regions work cohesively as neural population. In this manuscript, the Desikan Killiany atlas is used as a parcellation scheme as it has been used successfully before for whole brain simulations (Cabral 2011, Cabral 2012). Only the cortical areas without the insula are represented in the model constituting a total of 33 regions for each hemisphere for a total of 66 brain regions. These regions represent the nodes in the network model, and the network neighbours are defined using white matter tractography to map out fibers that run between two connected regions of interest. The change of the activity in the i^th^ brain region is defined as follows:

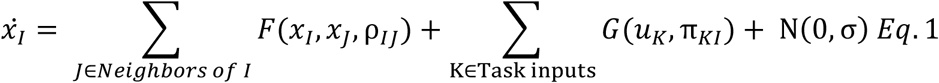

The first term represents the network component which is described by a function F that depends on its own activity x_i_, activity in its neighbour x_j_, and the physical properties of the fiber represented by the vector ρ (i.e., the number of fibers between regions, the delay in propagation). The second term consists of a function G that represents external input, whose activity is represented by a k-dimensional vector u representing all sub-cortical and sensory inputs, and the vector π representing again the physical properties that project these inputs into the cortical model (i.e., thalamic tracts into cortex). The last term represents a zero-mean Gaussian noise from the neuronal populations or from omitted higher order terms from the network equations. For resting state activity, the assumption is that *u*_*k*_ (t) = 0 ∀ t and the first term dominates the change in activity. This still leaves a large family of functions that are used to approximate F, with many parameters that can widely change the dynamics of the system. In theory, all of these functions can be used to fit the fMRI data with the Neural ODE algorithm, and for each of them initial conditions can be estimated from empirical data. In this manuscript, we focus on the simplest model the Firing Rate Model and solve for its parameters as well as for the noise level in the system.

#### 3.2.1 Firing Rate Model

The FRM represents the activity of a brain region as the mean firing rate. The change in firing rate of a region is a weighted sum of all its neighbours’ activity (Eq 1). The FRM has two parameters: the global coupling parameter k, which controls the strength of network input, and the level of noise amplitude σ, which simulates random activations of brain regions due to unknown neuronal activity (Cabral et al., 2012). At values of k< 1/max eigenvalue of W, the system is stable and the system decays to the origin without extraneous noise input. Typical values are k=0.9/(max eigenvalue) and σ=0.3 (based on the Desikan Killiany atlas) where there is a trade-off injecting noise in order to perturb the dynamics and the relative strength of the network to keep the neural areas functionally linked over time (Cabral et al., 2012).

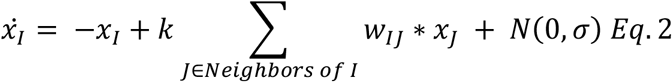

#### 3.2.2 Parameter and Structural Perturbations

Each of the components of the FRM are varied, the global coupling parameter k, the amplitude of the noise level σ, and the structural weight matrix W. The global coupling parameters and the structural matrix are fixed before using the Neural ODE algorithm, such that for each specific set of values for k and W a separate LSTM network is trained to generate the initial conditions. For the global coupling parameter variation, the parameter k in Eq 1, is varied from 0.9 to 0, in 0.15 intervals. To generate the structural perturbations, a random percentage of the original edges are swapped, to connect two different nodes, while keeping the graph symmetric. In this manner, random perturbations are created starting with the original structural matrix and randomizing the location of the edges. Each of these graphs would result in different dynamics, but the trajectories from the model containing the original structural connectivity should be the closest to the measured rs-fMRI data. This is varied by swapping 90 percent to 10 percent of the original edges, and expecting a variation in the accuracy in the models between the random matrices vs the original structural connectivity. The noise parameter is not used in the LSTM Neural ODE calculation, as they are integrated during training without noise. However, during testing after the initial conditions are estimated the noise is introduced by testing σ using values [0.0001, 0.15, 0.3, 0.45].

#### 3.2.3 Null Models

In order to compare the effects of fitting to the Neural ODE with other functions, the rs-fMRI data is fit with null models that are not simulating network activity. The simplest of these models, sets the Neural ODE equations to 0, such that the prediction of the LSTM is the output of the model. This quantifies how well the LSTM network’s initial condition prediction matches the next predicted output without any of the BNM functions. For future timesteps, it acts as a simplified autoregressive model by holding the current input as the output, i.e. x(n+1) = x(n). This is implemented by setting the connectivity matrix in Eq 2 to an identity matrix which cancels out with the first term and sets the equation to 0, and is referend to as the Autoregressive (AR) model. The second null model is obtained by setting the global coupling parameter to zero in the FRM and the differential equations reduce to an exponential decay. These models should perform worse than the BNM equations but test the limits of the global coupling values. Finally, we compare it to a pure Machine Learning Inference model as published in Kashyap & Keilholz 2020, where at each timestep the output of the LSTM is fed in as the next input. This model is non-deterministic as the output of the LSTM is sampled from a distribution. The function implemented by the LSTM in this case is completely unknown, as even the noise level changes as a function of the input. This model, however, gives a good estimate of an upper-bound on predictability on a short time window and does better than using traditional BNMs and can replicate complex resting state processes as Quasi Periodic Patterns and dynamic functional connectivity analysis performed using K-means (Kashyap & Keilholz 2020).

### 3.3 Neural Ordinary Differential Equations

The Neural ODE algorithm estimates the initial conditions of a given dynamical system based on previous observations. An overview of the algorithm is shown in Figure 1, based on using Neural ODE to fit a spiral dynamical system based on noisy observations. The task of the recurrent neural network is to predict the true initial conditions of the spiral dataset (shown as blue underlying trajectory in Figure 1) based on the sequence of observed measurements shown in green. A RNN implementation known as Long Short Term Memory (LSTM) was used to perform this task, as it keeps the information of past data observations [x_0_ to x_t-1_] in its hidden state p_t-1_ (Graves & Schmidhuber, 2009). Thus, when the timeseries is fed into the RNN one timepoint at a time, the current information is incorporated into the hidden state and is passed forward as shown in the LSTM unrolled version, in order to aid in the prediction of future observations. As the LSTM observes more datapoints, its predictions become more accurate up to a certain limit, after which newer data does not add any more information to what is already contained in the hidden state (Graves & Schmidhuber, 2009). For this particular task, the LSTM is trained to predict the initial conditions of a given dynamical system based on the observed timeseries. Since the initial conditions are not known and thus an effective gradient cannot be computed based on initial conditions alone, the algorithm assumes that the next observation is the integral of the predicted initial conditions and the given dynamical system with some noise added to it (Chen et al., 2018). Thus, the loss function is calculated with respect to the measurement at the next timestep in order for the output of the LSTM to converge onto the correct initial conditions for the given timepoint.

**Figure 1.**
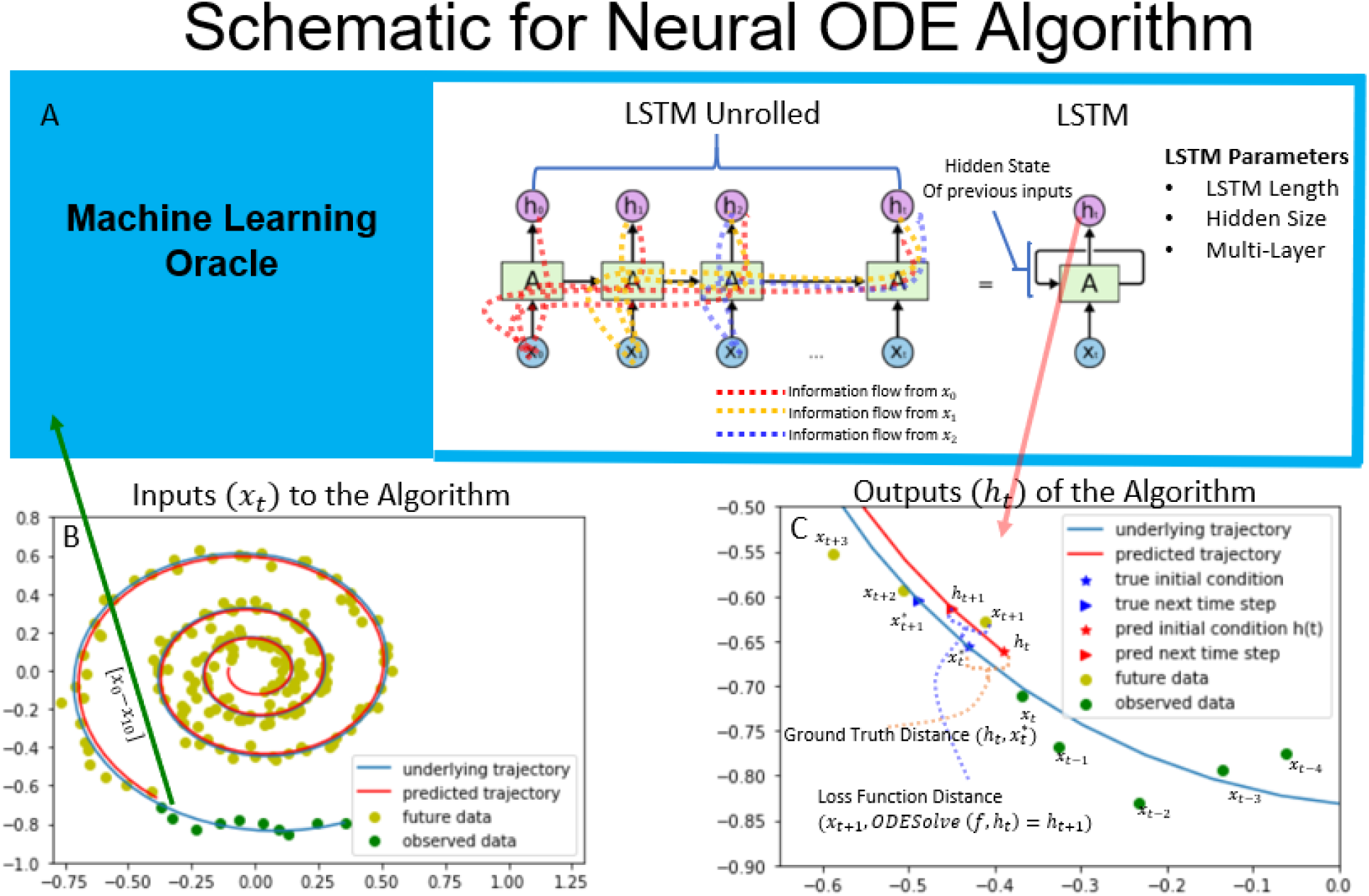
Schematic of the Neural ODE Algorithm. Schematic of the Neural ODE algorithm. An example spiral trajectory is shown in panel B. Sequences of the data (the green points) are fed into the RNN which updates its hidden state with more input values. The RNN sees one data point at a time and updates its hidden state as well as outputs its prediction for where it believes the initial condition to be. Information from previous observations is kept track of using the hidden state that is carried forward to future timesteps. An illustration of this concept is shown using the RNN unrolled diagram. The output of the RNN represents the initial condition of the dynamical system at that timestep. For the spiral dataset, the true ground truth initial condition 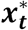 is known and is illustrated in bottom right. The ground truth distance is used to determine if the trained RNN network can successfully predict the initial conditions in the constructed dataset. The next timepoint is predicted by integrating the ODE system based on the given dynamical system. The loss function is defined as the difference between the next predicted timepoint and the next observed time point, x_t+1_. Minimizing the loss function distance minimizes the ground truth distance to the initial conditions.

The following steps outline the sequence of execution used to train the system.

1. The input data is shown in the bottom left corner Figure 1, where it consists of sequence of timeseries x_0_, x_1_, x_2_… to x_t_. They are fed into the LSTM network one at a time. The Tensorflow LSTM implementation of the RNN remembers up to x_t-N_ values where N is the length of the LSTM. N is set sufficiently large such that the performance of the LSTMs converges to its optimal value in M timesteps and N>M (see supplementary section 3).
2. For the current input x_t_ and the LSTM with its hidden state p_t-1_, the LSTM estimates the initial conditions as a normal distribution with a mean and standard deviation (μ_t_,σ_t_). We then sample an estimate of the initial condition h_t_ from this distribution. This is not explicitly shown in Figure 3.1 but is done in order to avoid overfitting.
3. Calculate the loss function at point t (Figure 3.1 bottom right): Using a 4th order Runge-Kutta to compute ODESolve(f,h_t_), where ODEsolve represents the differential equation solver to integrate over the interval t to t+1. This point is assumed to be close to the next measured datapoint x_t+1_. The Euclidean Distance between these points is used as the loss function. A graphic of this distance is shown in Figure 3.1 bottom right.
4. The gradient is then calculated with by averaging the loss function across all timepoints as well as across all batches and is backpropagated through the Neural ODE using Tensorflow, in order to train the network.

The algorithm trains until the loss function distance approaches zero. In practice, this is implemented by training to X number of steps and X is varied to make sure that the algorithm produces consistent results (supplementary section 4). The Neural ODE algorithm assumes that the ground truth distance between the predicted initial conditions and the true initial conditions also converges to zero when the loss function distance is minimized. The algorithm is initialized with a null hidden state, and slowly updates its hidden state as it sees more observations over time.

### 3.4 Tensorflow Implementation

The schematic above is implemented in the architecture shown in the Figure 2, which utilizes parallelization to efficiently train with large amounts of data. The network trains simultaneously using multiple subjects (batching of 80 subjects) and multiple time steps at the same time (50 consecutive time steps in Figure 2 but varied in supplementary section 4). The input would be a vector from the activity of all brain regions (66 ROIs at one time point) and is 3-dimensional tensor (66 ROI regions, 80 batches, 50 timepoints) and hidden state vector calculated from the previous time step by the LSTM. The LSTM is implemented using the CUDA library for NVIDIA GPUs and the architecture is a 4-layer hierarchical LSTM network. The LSTM is first initialized with a hidden state of all zeros and computes the data sequentially starting with the first timepoint of the 50 timepoints. The hidden state of the first input timepoint is then passed to the next timepoint, and the last state is passed as an input to the next time segment. The hidden state represents the feedback arm of the LSTM and is set to 150 (varied in supplementary section 4) and is 4 layers deep. The final output of the LSTM is then transformed from the size of the LSTM (150) using a feed forward Neural Net to the size of the initial conditions before it is fed to the BNM. The initial conditions are represented as a normal distribution with a (μ,σ) and the initial conditions is sampled from that distribution. The final output represents the mean firing rate of the neural population at that timestep for the Firing Rate Brain Network model (66 for the different brain regions). The model is then integrated for a time interval using a 4^th^ order Runge Kutta ODE solver using the Firing Rate BNM and produces an output representing the activity of the measured signal at the next time signal. The gradient is then subsequently calculated by taking the difference of the predicted output and the next measured timestep and then used to adjust the weights of the neural networks such that the sampled output of the LSTM converges to the initial conditions of the BNM. Since the accuracy is a function of the size of the network with respect to the data size, the parameters for the neural network were determined via sweeping each of these parameters across several values. The details of this process are outlined in supplementary section 4.

**Figure 2.**
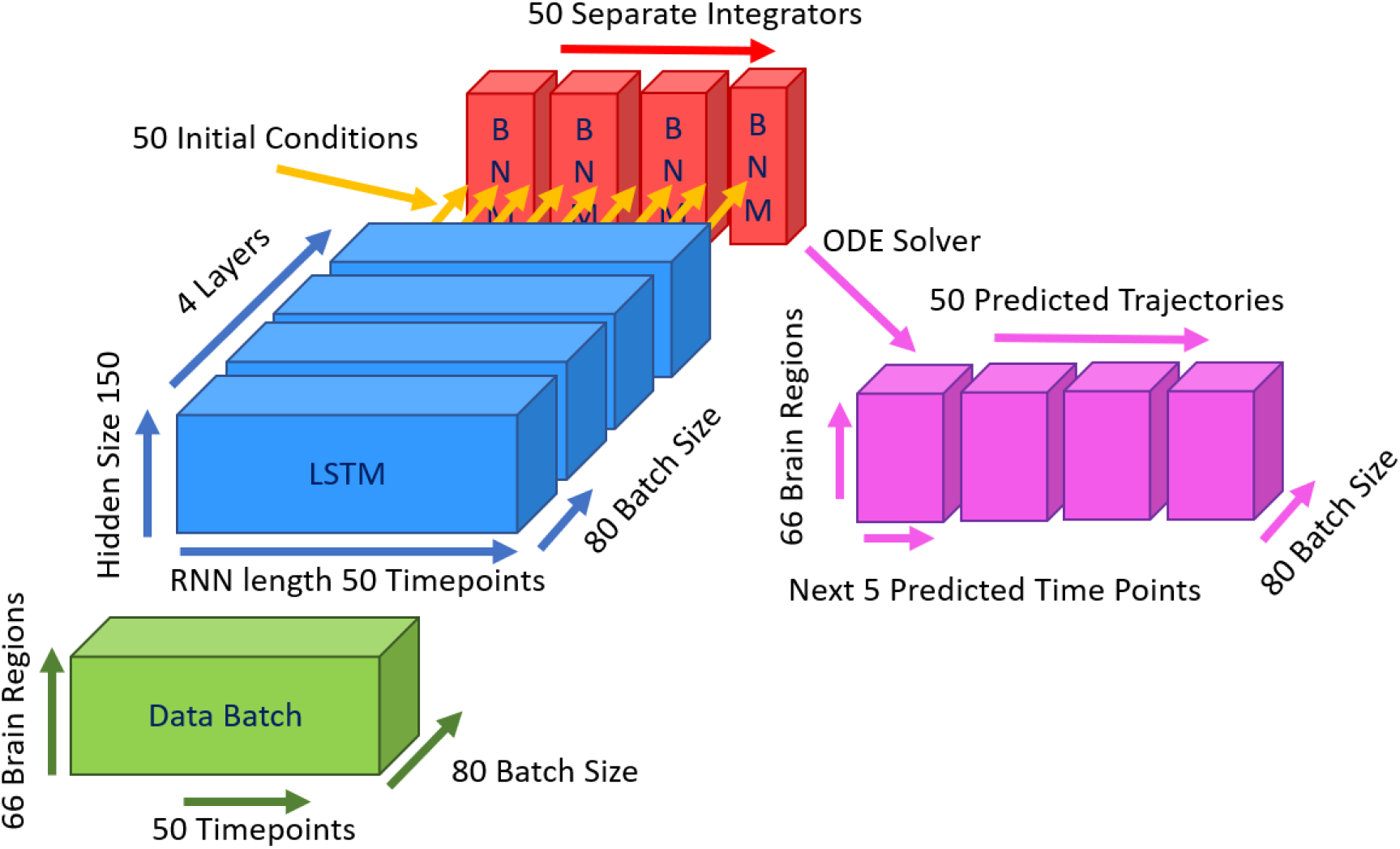
Neural ODE Architecture. Neural ODE architecture as represented by tensors with their appropriate sizes. The Data Batch consists of 50 consecutive timepoints from 66 Brain Regions of 80 subjects (batch size). The batch is fed into a 4-layer LSTM network, with a hidden size of 150. The initial conditions of the LSTM are set first to zero, and then to the output of the previous batch that contained the last 50 timepoints. The output of the LSTM is transformed via a feedforward layer to represent the mean and standard deviation of the initial conditions. An initial condition for each timepoint is then sampled and fed into the BNM in parallel and integrated by 50 separate integrators for the next 5 consecutive timepoints. The prediction at the next timestep is used to train the system, while the other 4 timepoints are evaluated during testing.

**Figure 0.**
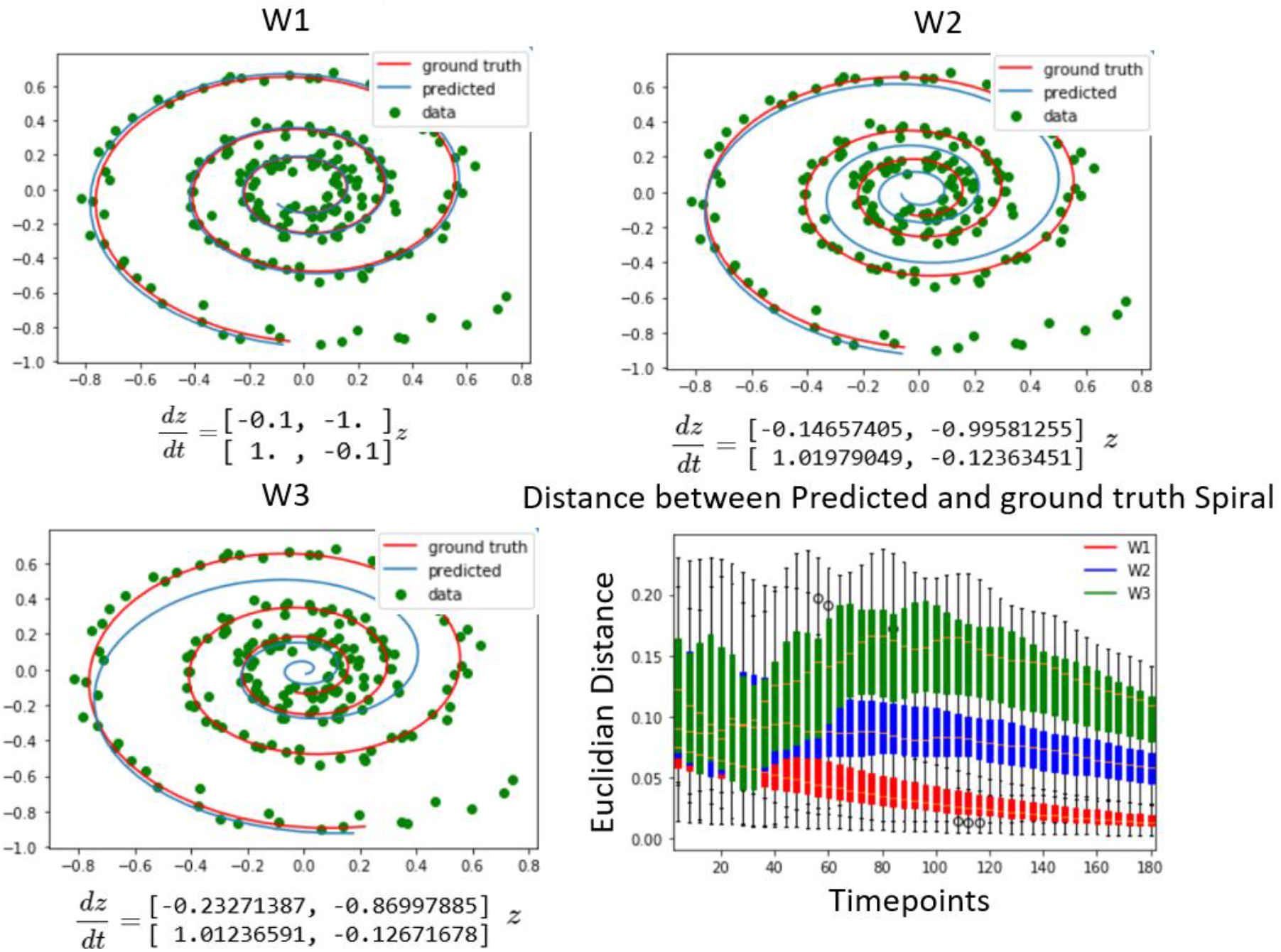
Differentiating between Dynamical Systems on Spiral Data. Using a candidate spiral in the neural ODE algorithm that is a noisy version of the ground truth spiral. The top left shows the alignment of the data, while fitting the spiral with a weight matrix identical to what was used to generate the data. The top right and bottom left figures show the alignment with spirals increasingly further away from the ground truth. The neural ODE algorithm is able to fit any of the spirals to the data for a given set of observations, but over time the candidate spirals that are further away from the ground truth diverge much faster from the future data points. This is quantified by the plot on the bottom right where the distribution of the three different spirals distance to the data is plotted from the predicted initial conditions. The distance between the spirals of different weights and the observed data points is initially close but diverges further away when compared to future timepoints. Note due to all trajectories going towards the origin, the distance at very large timescales converges to zero, but this is not expected in the brain data, where the signal does not approach a single attractor.

During testing, the trained LSTM model is used in the inference mode to estimate the initial conditions at each timepoint; the gradient is not computed at this stage. From each of these initial conditions x_init_ estimated from data observation x(t) future timepoints x(t+1), x(t+2), x(t+3) are generated by integrating the according to the equations of the Brain Network Model. In this manner, estimates at the same sampling period as the data are produced and the resulting trajectories are compared with the measured trajectories. To quantify the distance between them, the adjusted r-squared is taken at each future timestep between the vector representing the simulated activity of all 66 regions and the measured mean fMRI activity in the corresponding regions.

### 3.5 Experimental Data

#### 3.5.1 Structural Network for Brain Network Model

To estimate the structural network, tractography was run on 5 HCP Diffusion Weighted Images using the freely available software Matrix (Kashyap & Keilholz, 2019; Van Essen et al., 2013). The fiber orientations in the DWI images were first estimated using constrained spherical deconvolution. Then, using a probabilistic streamline algorithm, 100 million fibers set at a maximum length of 250 mm were computed for each individual and then filtered to 10 million fibers. To construct the structural network, we estimated the number of fibers that intersected two ROIs in the Desikan-Killiany atlas and normalized the power by dividing by the surface area of the receiving region (Cabral et al., 2011; Desikan et al., 2006). The matrix is finally normalized by dividing by the largest eigenvalue such that the graph Laplacian (k*SN-I) has only negative eigenvalues (Cabral et al., 2012). This normalizes the dynamics so that the feedback decays over time and does not exponentially increase the signal over time.

#### 3.5.2 fMRI Data

The fMRI data was obtained from the Human Connectome Project 447 Young Adult subjects release were used as functional data to test and train the models. The scans that were downloaded were pre-registered to Montreal Neurological Institute (MNI) space in surface format (MSMAII). The scans were first denoised using 300 Independent Component Analysis that are provided from HCP using the steps they recommended (Salimi-Khorshidi 2014). The surface-vertex or grayordinates time series were transformed to the ROI time series by averaging all vertices according to the parcellations established by the Desikan-Killiany atlas. This was done on an individual level since the surface parcellations are provided to by HCP and Freesurfer for each individual subject (aparc and aprac2009 files). The signal is then bandpass filtered from 0.0008 Hz to 0.125 Hz and then the global signal regressed using a general linear model using the mean timeseries of all cortical parcels as the global signal. The final signal is subsequently z-scored (Kashyap & Keilholz, 2019). For the task data, each dataset was processed separately (language, working memory, motor, social, emotional, gambling, relational) and then concatenated together. Each task dataset was truncated to the closest multiple of 50 timepoints and the autoencoder is fed alternating segments of task and the rest data for training. We trained our algorithm using task data as well as rest data, because the algorithm was able to perform better on most metrics when trained with more varied data. Moreover, it is thought that even during task activity, resting state networks dominate most of the cortical activity and task networks often look indistinguishable from rs-fMRI networks (Smith et al., 2009). However, during evaluation we have only shown our results on predicting future resting state fMRI and are planning to address task in our future work.

### 3.6 Metrics and Evaluation between Simulated and Empirical Trajectories

The dynamical models are evaluated on how well they fit with the empirical observations from the estimated initial conditions. For the spiral dataset, the initial conditions are already known and therefore, the results can be quantified directly using a Euclidean distance between the estimated and true initial conditions. To quantify how well they fit for the fMRI data, the r-squared and the mean squared error at each timepoint was calculated between the vector representing the activity of the 66 brain regions of the predicted and the observed data. Since the loss function of the Neural ODE algorithm, converges to zero during training across most models, this metric tends to be most similar when comparing across models (see supplementary section 5). Therefore, in order to differentiate between the models, the error is calculated for subsequent timepoints to gauge how well the trajectory follows the timeseries over a longer period of time. The results are calculated across a batch of subjects (N=80) randomly selected from the dataset and are calculated for each initial condition estimate at every timepoint. The difference between group and individual variances well as the effect of testing at each timepoint vs only a few timepoints is shown in supplementary section 5. While there is no pronounced difference in testing a few timepoints and evaluating the system at every timepoint, there is a large difference in variance between the averaging the error out in a batch of subjects (N=80) vs in the individual models. The group metric is selected as it has a smaller variance and is more robust, and the application of Neural ODE is to fit a group model rather than an individual model for the HCP resting state dataset.

## 4. Results

The goal of this work is to determine if the Neural ODE technique can be used to solve for the initial conditions of BNMs and then, based on simulated trajectories generated using these conditions, differentiate between different models based on how well it follows the consequent rs-fMRI measurement. To demonstrate the validity of the approach, the methodology is applied on a constructed spiral dataset where the true underlying dynamical system is shown that the correct coefficients are determined using this approach. Then the algorithm is applied using the FRM to fit with the rs-fMRI dataset and determine its parameters and coefficients.

### 4.1 Differentiating Between Dynamical systems on Spiral Data

The generation of the spiral dataset is given in detail in supplementary section 1-3, but, in essence, it is a two variable linear dynamical system that is often used as an example in machine learning literature to show the feasibility of solving for initial conditions. The supplementary sections first demonstrate the results from the previous paper Chen et. al 2019, where the Neural ODE algorithm’s predictions are able to converge to the right initial conditions after observing a sufficient number of previous timesteps in order for the LSTM to make valid predictions. The following sections then show the methodology of finding parameters of the network such as the hidden size and number of layers. Larger networks are more sample-efficient as they can predict the initial conditions based on fewer timepoints, but the overall accuracy has a limit. At a certain size, the accuracy does not change much with alterations to the network and is used as the model to perform the following experiment on system identification.

The spiral data was generated by a known system of differential equations as shown in Figure 3 top right. The purpose of the Neural ODE algorithm is to solve for each of the candidate dynamical systems the initial conditions with respect to the observed data. Each of the systems contain different coefficients in their respective weight matrices, where the W1 represents the original dynamical system and W2, and W3 contain the original structural matrix perturbed with increasing noise. The results in Figure 3 show three examples of fitting these spirals to generated spiral data for different weight matrices. In the examples, the spiral data used to generate the data, W1, follows the future trajectories the closest, but for the first few points after the initial conditions the other candidate systems do quite well in fitting to the data and sometimes even slightly better than the original system. Figure 3 (bottom right) quantifies the distance between the predicted and observed trajectories over many examples. The spiral matrix W1 is closest in Euclidean distance to the data over long periods of time, but the difference is less pronounced at short time periods where the distributions overlap with W2 and W3. Therefore, since the Neural ODE fits any dynamical system tangentially in time, it is important to observe the long-time dynamics to differentiate between the models. At very long intervals, the distance starts to decrease as all trajectories converge to an attractor based at the origin, which is special for the spiral dynamical system and not present in the neural data. In summary, the results on the synthetic spiral data show that it is possible to use this method as a system identification, but the systems need to be simulated for a long enough time interval for the differences to manifest, and the distances close to the initial condition are harder to tell apart, since the output of the LSTM minimizes the prediction error at the first timestep.

### 4.2 Fitting differently parameterized Firing Rate Models to resting state fMRI data

Most BNMs have many parameters that affect the dynamics differently and can be tuned to individual fMRI data. Therefore, it is essential to test whether this method allows us to differentiate parameterized BNMs. The parameters of the FRM have been estimated in the past, and the method can be validated by reproducing previous estimates (Cabral et al., 2012).

In Figure 4, differently parameterized FRM models as well as three Machine Learning Null models are compared in their ability to reproduce the future trajectory. After estimating the initial conditions, the distribution of distances between the predicted and the real trajectory are plotted over time. At the first timestep (Figure 4, top left), all the models perform relatively similarly as a direct result of minimizing the loss function using an LSTM. Similar to the spiral example, the models perform differently when moving forward in time. At the fourth timestep in Figure 4 (top right), surprisingly the exponentially decaying null models with a zero global coupling, and the autoregressive null model with no BNM (labeled as AR) perform better than network containing the brain network equation. This suggests that the introduction of any BNM, decreases the accuracy of the model and the model performs best using just the LSTM predictions. The null model utilizing LSTM inference (Kashyap & Keilholz 2020), performs the best and represents an estimate of an upper bound in predictability of the rs-fMRI signal.

**Figure 4.**
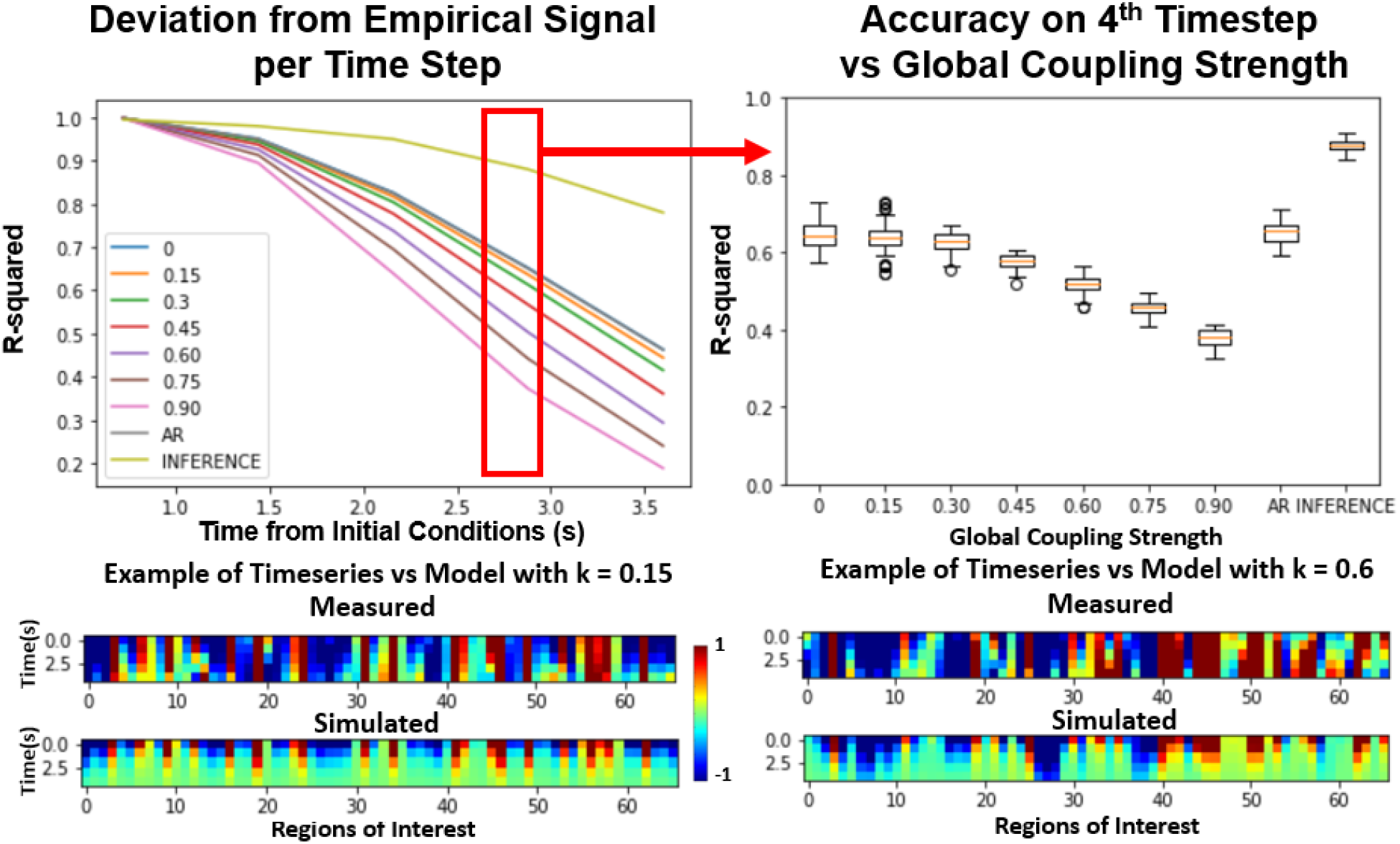
Effects of Global Coupling in the Noiseless Firing Rate Model. Evaluating the FRM (Eq 1) with different global coupling parameters. Top Left: Error per timestep from the estimated initial conditions for various different parameters compared. The y axis represents the r-squared value, between the simulation and the model. Examples of the model timeseries vs the resting state timeseries are given on the bottom at two different parameterizations but the distribution shown in the top panels quantify their performance across 2500 trials. Top Right: At the fourth timestep, the distribution of the r-squared across all the models is plotted. For the FRMs, the accuracy decreases with the increase of global coupling, and the model with zero global coupling performs the best. The Autoregressive (AR) model utilizes the LSTM for the first timestep prediction and then outputs the next prediction as the previous timestep and does as well as the FRMs with zero global coupling. The inference model using LSTM at every timestep as implemented in Kashyap & Keilholz 2020, performs the best in terms of accuracy, but the model dynamics are unknown as they are implemented using deep learning.

However, interestingly this trend completely reverses when noise is introduced into the models. In Figure 5, both the standard deviation of the noise as well as the global coupling parameter are varied. At low noise levels, models with low global coupling outperform the models with high global coupling. However, with increasing levels of noise, the BNMs with stronger network effect outperform the models with low levels of global coupling. This result shows the importance of noise in establishing the parameters of the FRM, as the properties of the structural network become more important when the system has high noise. The overall r-squared of the models also decrease with the introduction of noise, but the rate at which they diverge from the measured trajectories seems to be a function of the global coupling parameters. Previous FRMs that used the same brain parcellation simulated with k = 0.9 and σ = 0.3, and noise was seen as essential in simulating the BNMs (Cabral et al., 2012). However, since both parameters are unknown and the overall r-squared decreases with the introduction of noise, just based on varying these two parameters it is difficult to conclude which parameterization yields the best result using this approach. The AR null model still performs better than the introduction of a BNM, but the difference is much smaller than before. The inference model is not included in the noise estimations, as the dynamical system is represented as a RNN and cannot be manipulated in a controlled manner as the other models.

**Figure 5.**
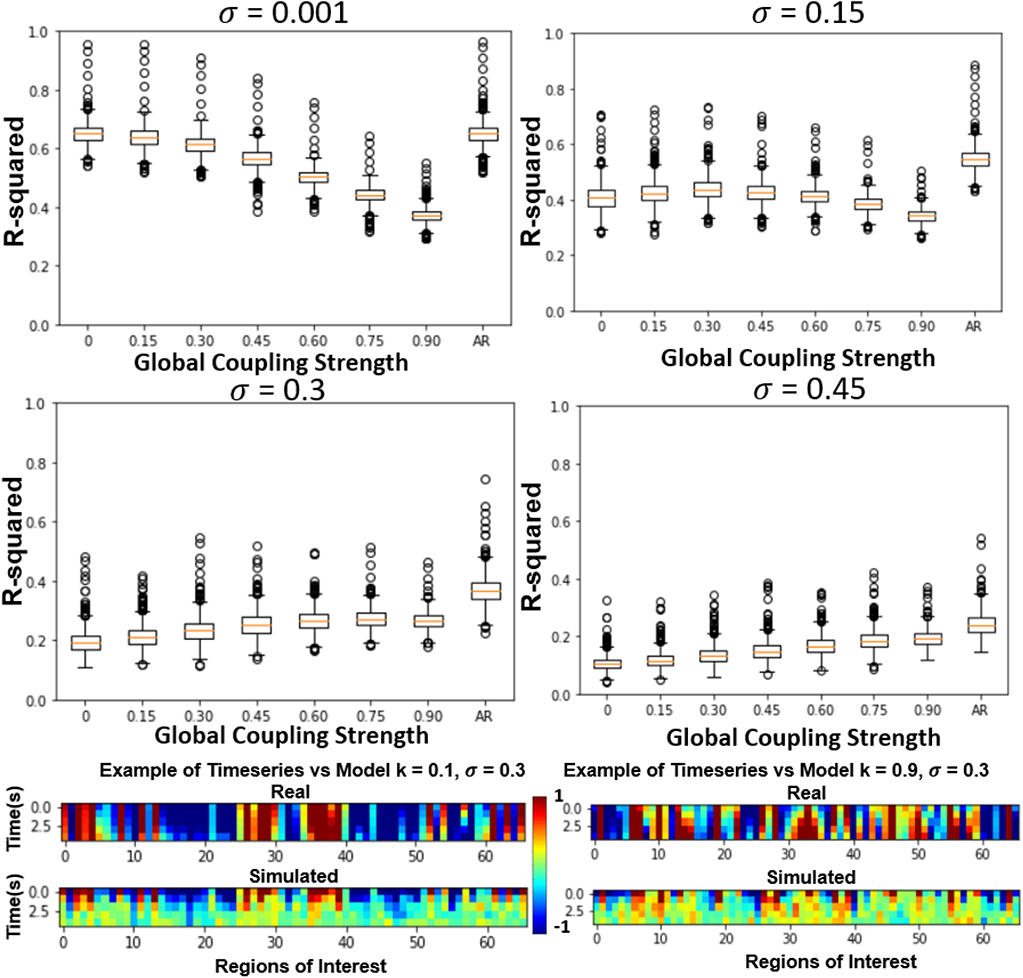
Effect of varying Global Coupling and Noise on the 4^th^ Timestep Accuracy of the Firing Rate Model. The effect of varying the two parameters in the FRM, the global coupling K and the standard deviation of the noise σ. The introduction of noise lowers the accuracy of all models but does so in an uneven fashion. At low levels of noise, the exponential only model (k=0) outperforms the BNMs with structural connectivity matrices. This trends reverses with the introduction of increasing noise power, where the BNM with stronger global connectivity matrices seem to outperform the naïve exponential models. The autoregressive model also performs worse with increasing noise levels, and although it still outperforms the FRMs regardless of the coupling strength, the gap between them is reduced.

The introduction of noise changes the resulting dynamics and can be also seen via the time traces. In Figure 5 bottom, the time series without noise result in trajectories that decay to zero. This follows a well-known analytical solution of the consensus equation, where the values of a connected network with eigenvalues less than 1 converge to the origin (Mesbahi & Egerstedt 2010). However, the introduction of noise results in more complex trajectories as shown in Figure 6 bottom, where the values do not decay to zero, but rather randomly oscillate around the origin which acts as an attractor in the system (Cabral et al., 2011). The role of the structural network, in this case becomes more important as it magnifies the noise inputs through the network, and at high noise levels the FRM with global connectivity evolves more similarly to the measured rs-fMRI signal.

**Figure 6.**
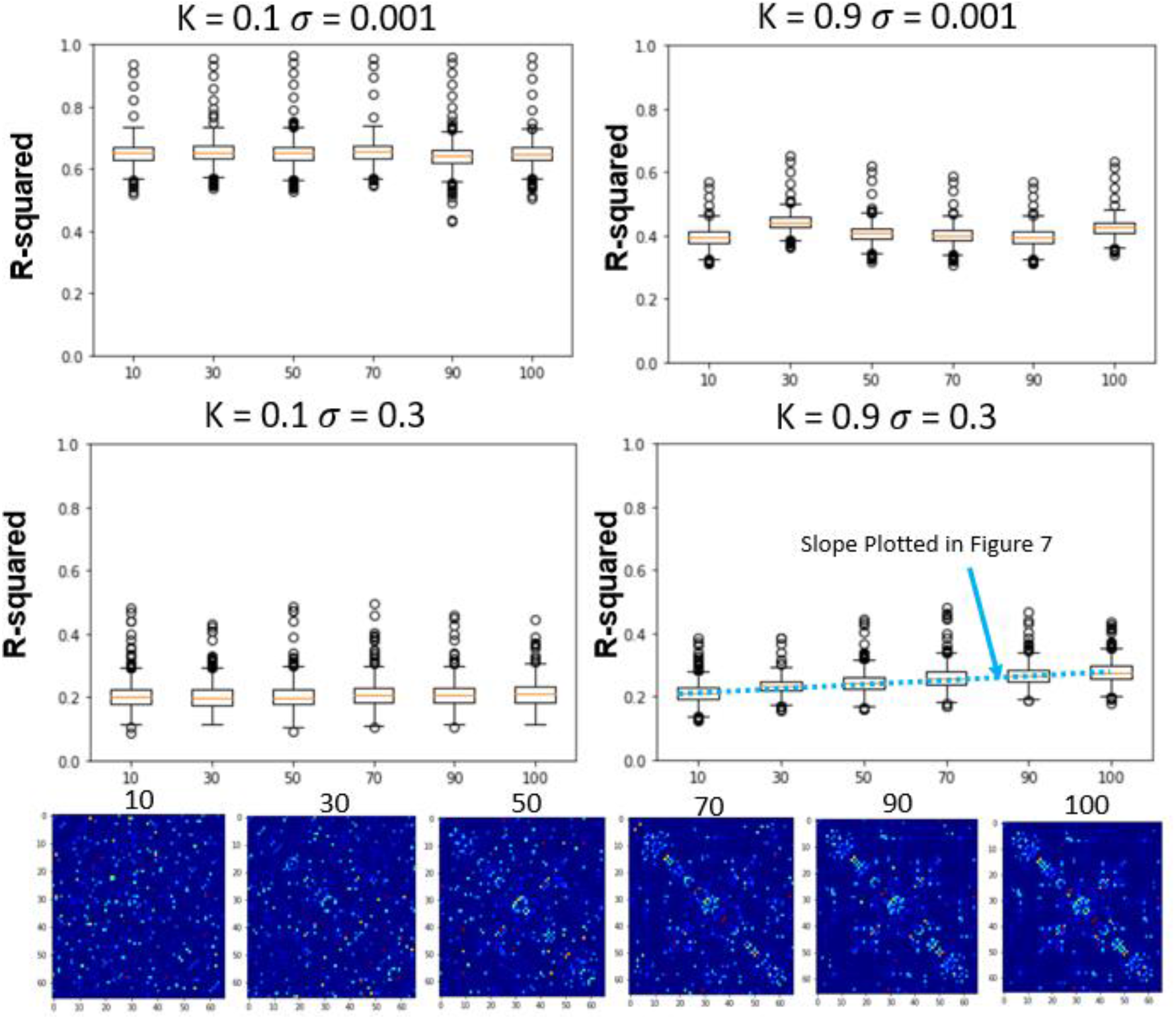
Effects of Global Coupling, Noise, and Structural Connectivity on the 4^th^ Timestep Accuracy of the Firing Rate Model. Examining the effects of parameterization and estimating the correct SC. At top left, we show the results of changing the structural connectivity for a low global coupling model. It does not vary as a function of the structure and performs relatively similarly to the LSTM only null model. At high global coupling, the models show that they are more of a function of the correct structural network, although they perform worse than at high global coupling. The slope across the performance of different structural connectivity (bottom right of Figure 7) is used as a metric in the next section to solve for global coupling and noise levels.

### 4.3 Differentiating Between BNM due to differences in Structural Connectivity

In the previous section, only the parameters of the FRM, the global coupling strength as well as the magnitude of the noise were changed. In this section, the effects of simulating six different SC matrices at high and low different global coupling (k=0.1, 0.9), are quantified while varying the high and low noise levels (σ=0.001,σ=0.3). The SC is varied from the measured SC by flipping edges and results in SC seen in Figure 6 bottom row. The r-squared value at the fourth timestep, between the different models is plotted in Figure 6 top two rows. Unlike the previous sections where the correct values of the parameterizations were unknown, in this experiment, the dynamical system with the original structural connectivity is expected to outperform the models with perturbed structural connectivity.

In the low noise levels (σ =0.001), the model shows no relation to changing the structural connectivity for either of the coupling strengths. At the high noise levels (σ =0.3), although the model has a lower r-squared than at the low noise levels, the effects of the network are observed, where the original SC outperforms the corrupted versions for both low and high coupling strengths. The trend is once again more pronounced for the high global coupling (k=0.9) when compared to low global coupling (k=0.1).

### 4.4 Estimating the parameterization and noise level of the Firing Rate Model

Since both the noise level and the parameterization of the system is unknown, just by looking at the r-squared between the simulated and measured systems as seen from the previous sections, its not possible to find the system that is most predictive of the data. However, in the previous section the system was more sensitive to the measured structural connectivity that certain global and noise levels. Therefore, using the slope calculated at different SC values, different noise levels and global coupling values are simulated, to see where the slope is maximized when changing the structural connectivity. We hypothesize that at the correct noise levels and global coupling value, the system will be most sensitive to the underlying structural connectivity. The slope is plotted from timesteps 2 to 5, across different global coupling values and noise steps in Figure 7. At timesteps close to the initial condition, higher levels of noise are needed to differentiate the systems sensitivity to the structural matrix. However, at higher timesteps, the opposite is true where higher noise perturbs the system too much and the overall r-squared drops so low that the models become indistinguishable to each other. Therefore, the fourth timestep is used where the max differentiation between models occurs regardless of the coupling values, where the trajectory is far enough from the effects of the LSTM fitting and close enough in time to test the predictability of the models. At this timestep, the maximum occurs at the values (k = 0.925, σ=0.35). This value is very close to what has been used to be simulate FRMs (k=0.9, σ=0.3) from previous publications (Cabral et. al 2012). Their approach of parameterization here has been reproduced in Figure 7 (right most panel), which calculates the FC of a 12 min simulation and correlates the FC with the FC of the empirical data. The maximum at [0.875, 0.3] is in good agreement with the short-term measures and the previous estimates.

**Figure 7.**
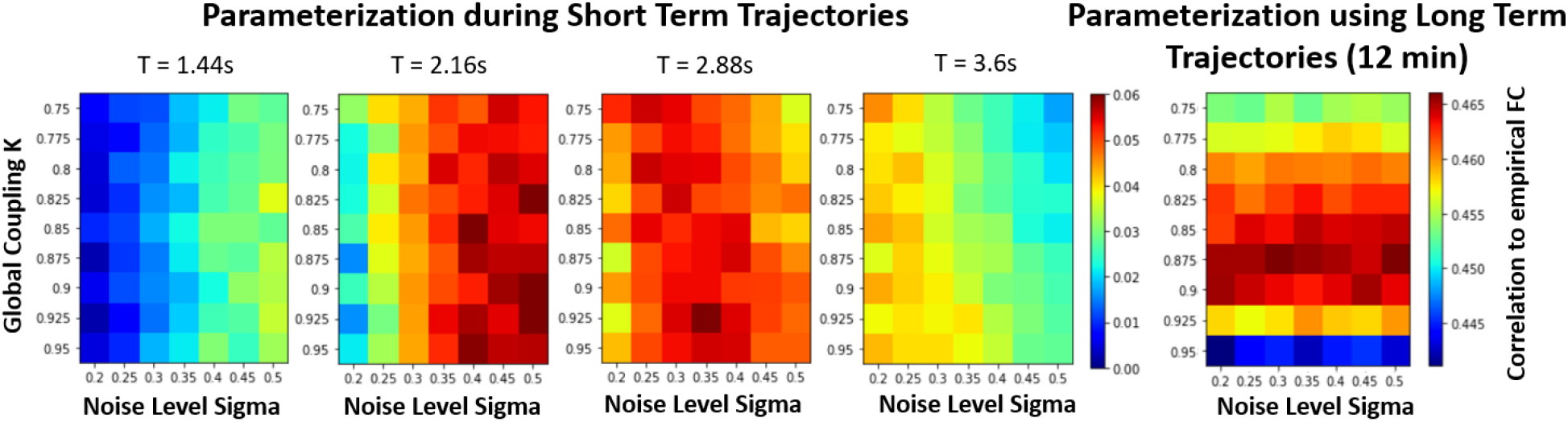
Parametrization of FRM using short term measures vs parameterization using Long Term measures. Examining the effects of changing the Structural Connectivity Matrices vs Accuracy (Slope in Figure 6) under different parameterization/noise levels of the Firing Rate Model. The performance to changes in SC is the only metric where the variation has an expected result; the original structural matrix performs better than a random matrix. The color bar represents the change in r-squared value and is plotted for different timesteps. At the fourth timestep (t=2.88) the maximum differences occurs on all the models regardless of the parameters, as at that time step the system is far enough to where all the models were synchronized in time using the initial conditions have sufficiently diverged and not too far where the overall r-squared drops so low that the systems are indistinguishable. At this timestep, a maximum occurs at (k=0.925, σ=0.35). Comparison to the traditional parameterization is shown in the right. Here the differently parameterized FRM models were simulated for 20 min and then the FC matrices of the resulting simulation were compared against the empirical FC. The traditional approach has a maximum at [0.875, 0.3], and previous reproductions of this experiment found a maximum at [0.9, 0.3] (Cabral et al., 2012). This shows a good agreement on which BNM recapitulates rs-fMRI in both short and long term measures.

## 5. Discussion

### 5.1 Overall Discussion and Significance

The study proposes using the Neural ODE technique to estimate initial conditions in different candidate BNMs with respect to measured data and then subsequently evaluating them on how well they evolve compared to the real data. In order to test this methodology, the technique was first applied to a well-studied spiral dataset to determine if our algorithm solved for the initial conditions when the ground truth was known and was shown that it could correctly identify parameters in a constructed example of system identification. Then the method was applied to a neural rs-fMRI data, to fit different Firing Rate BNMs using the Neural ODE technique by varying their 1) parameterizations, 2) noise level, and 3) by changing the structural connectivity. By using all three, the system was correctly able to identify the parameters of the FRM and is very close to previous estimations of the parameterization of the model using the same whole brain parcellation (Cabral et al., 2012).

The main results show that the Neural ODE is able to estimate for the initial conditions for a variety of different models ranging from simple exponential functions in our null model to the FRM that utilize the brain structure for its dynamics. In this manner, for the first time all these different models are able to be directly evaluated by comparing the predicted timeseries from the initial conditions with the empirical timeseries. The performance of first time step was roughly equivalent for all models since that was what the Neural ODE models were trained to predict, but the dynamics diverged from the measured timeseries when integrating the equations further out in time. However using this as a metric, the technique yielded a trivial model without any noise and a simple exponential decay function. Only under the presence noise, do trends emerge with respect to the global coupling parameter and the accuracy of the structural connectivity estimate. The model that was the most sensitive to changes in the structural connectivity input, also yielded a parameterization of global coupling and noise amplitude that matched previous results of the FRM (Cabral et al., 2012). Therefore, pertinent information of the BNM is present in short term trajectories analysis even though they are stochastic processes. The technique be extended to more complex BNM and be used to compare between them when the amplitude of noise is established. Unlike older metrics, this technique allows for direct timeseries comparisons between the theoretical models with measured experimental data, and circumvents the reliance on a certain metric/ interpretation of rs-fMRI. Moreover, it allows pathway forward in studying whole brain dynamics on a faster timescale, and illuminate what our current theoretical models can and cannot explain in terms of transient dynamics.

### 5.2 Parameterization and Noise Levels of the Firing Rate Model

The FRM, is the simplest of BNM used to simulate rs-fMRI, and has only two variable parameters in the model (Eq. 1), the magnitude of global coupling and the level of noise. From previous literature, the global coupling value of a traditional FRM is set slightly less than 1, around 0.9, which is just before the system becomes unstable. Closer towards k=0, the dynamics become an exponential decay according the Eq 1. The noise is usually set around 0.3 for this parcellation scheme (Kashyap et al., 2019, Cabral et al., 2012), where closer towards zero it turns the system to the well-known consensus problem where the timeseries converge to the attractor at the origin (Mesbahi & Egerstedt, 2010), and at higher degrees of noise the system becomes completely chaotic and non-deterministic. Searching for the correct parameterization of the FRM between the global coupling and the magnitude of the noise is therefore an important to simulate the model in the correct regime. Based on previous studies, the parameterization of FRM was estimated by maximizing FC in relation to the rs-fMRI FC, the parametrization was expected around (k = 0.9, σ = 0.3).

In practice, this relationship was not so easily ascertained by analysing the short-term trajectories as it was confounded by the presence/ absence of noise. The simulated trajectories that were the closest to the empirical trajectories were models that contained no noise. In the case of no noise, the FRMs that contained the structural network performed worse than the trivial exponential decay null models (Figure 4). However, in the presence of noise, the role of these network models seem to become important, as at higher global coupling the signal would deviate less from the empirical trajectories. The role of the structural connectivity here can be thought to averaging out the noise, and is more robust to local deviation due to noise introduced at each ROI. Supporting this argument, Figure 6 showed that in the presence of noise, the models also exhibited a dependence to changes in the structural connectivity, where the true structural connectivity resulted in dynamics closer to the empirical signal than noisy perturbations of the original structural connectivity. Although the exact value of noise cannot be solved by maximizing the r-squared accuracy while varying the noise amplitude as it results in a trivial result of favouring noiseless models; at the right noise/ global coupling parameterization, the accuracy of the FRM should be maximally dependent on the correct structural connectivity. This hypothesis was tested in Figure 7, where the change in accuracy of due to the changes in the structural connectivity was plotted. Using this metric, the FRM with (k=0.925, σ=0.35) is the most dependent to changes in the structural connectivity and is very close to previously known values that was being used for the FRM (k=0.9, σ=0.3) as well as our computed maximum using long term FC estimates (k=0.875, σ=0.3) (Cabral et al., 2012). The previous process used long term FC as a metric to maximize in order to parameterize the models, rather than using the short trajectories as in this manuscript, but here they show they give similar estimates on which FRM is closest to measured rs-fMRI dynamics. The evidence that these two values are close, suggests that our approximation of the true underlying dynamical system is at least scale free across the observed timeframes and models that have been used to simulate long periods of time, can capture meaningful dynamics in the shorter timeframe.

### 5.3 Comparison to other Neural ODE Architectures

The original Neural ODE implementation uses a backward time architecture, where the timeseries is inverted and fed into the RNN network, such that the first timepoint is fed into the RNN last and the final prediction is used to infer the initial condition of the whole timeseries and then integrated forward in order to compute the loss function (Chen et al., 2018). They do not evaluate the RNN prediction at every timepoint like in our implementation, but explicitly state that such an architecture would speed the training process. The Tensorflow RNN implementation page also recommended a parallel use of the RNN in order to speed up the training process (tensorflow.org/guide/keras/rnn). The innovative backwards time architectural method gets rid of the initialization problem of the RNN that exists in our forward time implementation but runs into a causality problem where future inputs influence the predictions of previous initial condition. Because BNMs are defined as a function of previous network activity, and because our intended use of the trained model is a continuous correction of the accompanying BNM model, the time forward architecture is used in order to solve for the initial conditions. The other significant difference is that the Github implementation of the Neural ODE also uses a LSTM after the ODE integration (Chen et al., 2018). This methodology is extensively evaluated in Kashyap & Keilholz 2020 but confounds our goal of comparing the fit of different dynamic systems, so it is simply presented as a null model labeled as inference in this paper. This model outperforms all other models in terms of short term prediction, but cannot be manipulated as in terms of the noise level, coupling strength or other meaningful biological variables. Rather it represents an estimate of an upper bound in terms of predictability seen in the rs-fMRI dataset.

### 5.4 Comparison to other techniques in Literature

The Neural ODE algorithm presented here is a relatively new technique first presented in 2018. To our knowledge this exact technique has not been applied in the context of fitting whole brain models with empirical rs-fMRI data. However, our methodology is quite similar to our own previously published work (Kashyap Keilholz 2020), but differs in the important following manners. In the previous paper, the system was trained in a very similar manner, but in the generation of new data from the initial conditions the older methods utilized the entire Machine Learning architecture, LSTM and the Brain Network Model to synthesize new data, whereas in this paper, the future timeseries is generated from initial conditions by integrating the Brain Network Model. The older method allowed to generate more realistic brain data and replicate brain dynamics better than traditional BNM as it utilized the LSTM in every timestep. However, this brought into unknowns into the dynamics and it was not possible to evaluate the BNM on their own. Therefore, in order to isolate the performance of BNM for the purposes of system identification, the LSTM was excluded from the inference process and was only used to generate initial conditions. A recent preprint (Wen 2020) also utilizes the Neural ODE approach to fit to rs-fMRI data. However, in that methodology it doesn’t use the Neural ODE tool to fit trajectories from the BNM rather analyzes latent variables of a model to predict task states. Many other approaches have started using different techniques for uncovering principles of dynamical systems in order to represent rs-fMRI (Basset 2020 paper, Singh et al., 2020, Hejm et al., 2017, Zalesky et al., 2021). Basset 2020 uses a similar r-squared metric to quantify the difference at the first time point prediction but does not extend this by predicting further out in time. In this paper, to our knowledge is the first to use these tools for comparing short term trajectories of given BNMs to measured rs-fMRI data.

### 5.5 Assumptions and Limitations

The error from the model’s prediction comes from multiple different sources such as 1) the mismatch between the differential equations and the actual dynamics, and 2) from the error in predicting the initial conditions, 3) inadequate descriptions of structural connectivity and/or the lack of including subcortical areas in the simulations.

1. A major limitation of this approach is to have an estimation of the underlying dynamical system that represents the data. This requires vast knowledge of what model including the specific parameterization might fit the dataset. However, the Neural ODE system is able to tangentially fit any dynamical system even trivial ones such as the exponential decay. Therefore, it is not really necessary to have a really good estimation of the underlying system and can be tested how well they predict subsequent timesteps.
2. We assume that for any assumed dynamical system the error from the RNN is uniform no matter what the function is, and the subsequent error calculated from the trajectories is due to the mismatch between the data and the dynamical system. However, this might not be true, and more complex models might have a larger errors in estimating initial conditions and therefore is a potential confounder in our analysis.
3. The inadequate description of the brain network is also a limitation and can be improved with higher resolution parcellations and subcortical areas. Another major drawback is the network is quite ill-defined because there is no consensus what constitutes a ‘cohesive’ neural population. However, different atlas and network definitions seem to give similar results suggesting that the principles of BNM are at least consistent across many parcellation schemes that are used today. However, results from previous literature, show that a more detailed description of the network only improves the models performance and its ability to recapitulate rs-fMRI and that coarser models are good enough for a proof of concept application.

Moreover, another major assumption and limitation of the approach is our choice of metric, r-squared used to compare the distance between two high dimensional vectors. It assumes that better models have a higher r-squared value, although they might be explaining trivial components of the signal. Other metrics such as derivative, or the relative phase between different regions of interest might prove as a much more useful metric to compare the predictions against the empirical signal. The method also introduces another variable on *when* to evaluate the differences of the model. Close to the initial conditions the trajectories are too close to differentiate, and as seen from the null models where the output of the LSTM already captures a large amount of the variance in the signal. Too far from the initial conditions yields trajectories that are too far away from empirical measurements and all models become completely indistinguishable. For our results, the fourth timestep (2.88s) was the most useful in differentiating between models, but this could vary from implementation and careful consideration needs to be used in interpreting the results and is a limitation in the approach.

### 5.6 Future Applications

The Neural ODE techniques has a lot of potential as an additional tool in conjunction with BNM. It can be used to evaluate any differential for brain data in real time by solving for the initial conditions. Moreover, it can be used to compare across increasingly disparate brain models that are being constructed for specific applications. For individual data, it seems especially promising, since the trained network can make predictions on an individual fMRI data and thus parameters of the BNM as well as the structural connectivity can be adjusted on the individual level.

For future approaches on more complex BNMs, it might be easier to assume the noise level and then the parameters can be solved in a more straightforward manner. The noise level seems to be endemic in the system, and rather than a parameter of the model. Since our mean surface area parcel is 858 mm^2^ according to our atlas, we estimate the cortical noise per area to be *N*(*μ* = 0, *σ* = 0.35 * *Sur face Area*/858 *mm*^2^) and not to be a function of BNM. Once the noise level has been established, the structural perturbations are not necessary and the coefficients of the BNM can be determined directly by comparing the r-squared of the models with the empirical signal. In this manner many more complex BNMs can be compared against each other.

## 6. Conclusion

This manuscript investigated whether by solving for the initial conditions of a Brain Network Model for a given observation of rs-fMRI data using Neural ODE, the estimated BNM trajectories based on these initial conditions would serve as a metric to differentiate between BNMs and the measured rs-fMRI timeseries. The approach used several different FRM to fit to the rs-fMRI data by varying the global coupling, noise, and structural connectivity. The results show that the parameterization of global coupling and noise that maximizes the model’s sensitivity to the structural connectivity, yields a model comparable to earlier parameterizations of the FRM. Therefore, the Neural ODE tool has the potential to differentiate and develop more complex BNMs to fit rs-fMRI data and a path to train the parameters on individual fMRI data.

## Supporting information

Supplementary Sections

