## Supplementary Sections for "A DEEP LEARNING APPROACH TO ESTIMATING INITIAL CONDITIONS OF BRAIN NETWORK MODELS IN REFERENCE TO MEASURED FMRI DATA"

### Supplemental Material

1. *Simulated Spiral Data*

In order to validate our approach, we tested our algorithm/architecture that we use on fMRI data, on a toy spiral data set. This spiral data has been used in the original Neural ODE paper (Chen et al., 2018), as well as in subsequent papers in order to test the validity of solving for the underlying dynamical system from noisy data. Our approach is to establish that our algorithm can reproduce the performance of the Neural ODE algorithm (Chen et al., 2018) on the spiral data and this therefore justifies its use on fMRI data. In Figure 1, we show how we generate our spiral dataset, namely by integrating a set of coupled differential equations with two state variables (equation shown in the left panel). The phase portrait of the dynamical system is shown, where all the trajectories from any initial condition spiral inwards towards the origin (Figure 1 middle). Time is not shown in the graph but is implied, where the first timepoint is on the edge of the spiral and the last one is the one closest to the origin. The derivative is large on the outside of the spiral and then decreases as it approaches the origin. Gaussian noise is added to the integrated trajectory to simulate measurement noise set at sigma = 0.03 (Figure 1 right). The goal of the Neural ODE algorithm is to estimate the underlying trajectory from the noisy observations that generated the data. For our experiments, we use the trajectories shown in the middle panel of Figure 3.3 as the ground truth and take the Euclidean distance between this trajectory and the predicted initial conditions as a measure on how well our algorithm performs.

**Figure 1 Generation of Spiral Dummy Data**

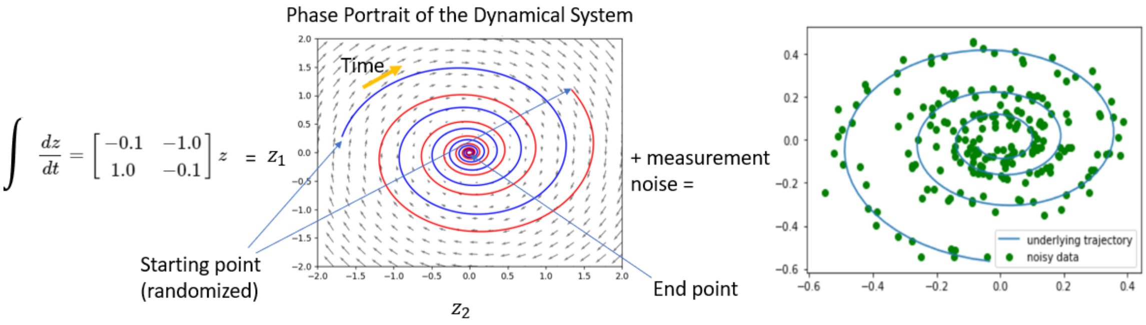


**The equation on the far left is integrated and produces the dynamical system shown in the phase portrait (middle). From two random initial conditions the red and blue trajectories spiral towards the origin, getting slower and slower as they approach the origin. The arrows indicate the magnitude of the gradient. We generated 1000 spirals from different initial conditions and added noise to simulate a noisy measurement. This results in the plot on the far right where the green dots represent the data that is fed into the algorithm. The goal of the algorithm is to be able to estimate the underlying trajectory shown in blue (far right) from the noisy observations. This will serve as our ground truth to check our predictions against.**

2. *Validation of the Neural ODE Algorithm on Spiral Data*

The spiral data allowed us to test how well our algorithm estimates the initial conditions, in a simplified situation where the initial conditions are known. In Figure 2, we show that the predictions of a trained network converge towards the ground truth of the initial conditions for the sample spiral dataset. Starting from the first observation, at the bottom right corner of the spiral, the LSTM makes a prediction based on all the observations it has seen so far, and then a trajectory is produced via integration of the dynamical system of the spiral dataset. The trajectories after seeing 7, 9, and 11 datapoints are shown on the spiral itself. The distance between ground truth initial conditions is also quantified to the right as a function of number of points the LSTM observes. The accuracy during the first couple timesteps is low because the hidden state of the LSTM is not initialized properly, but after enough datapoints are included, it is able to predict the initial conditions within a reasonable margin of error.

**Figure 2 Infer Initial condition for each Timestep on Spiral Data**

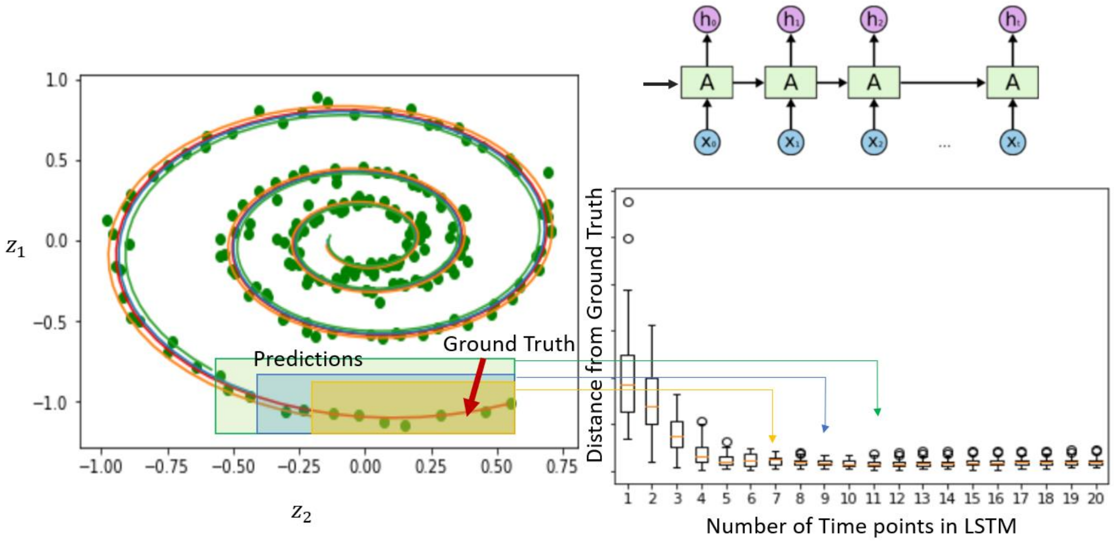


**Estimating the initial conditions for the spiral data. The figure on the left shows an example of a trained network estimating the initial conditions using 7, 9, and 11 timepoints. Each timepoint is fed into the LSTM sequentially (shown top right) one datapoint at a time and outputs the initial condition for that timepoint. The trajectories are then integrated and then compared with the ground truth (shown in red). The Euclidean distance between the predicted and the true trajectory is shown in right for a distribution of 40 unseen spirals. The accuracy converges after the first few timepoints during which the LSTM has not yet been initialized properly. After the LSTM has seen enough examples, the estimates converge slowly towards the true spiral trajectory.**

3. *RNN Parameter Estimations on Spiral Data*

Next, we test to see how our ground truth distance changes as a function of network size and parameters (Figure 3). For our given architecture, we can vary the number of layers (network depth), the size of the hidden layers, and the length of the LSTM network. The length of the LSTM network is a parameter due to Tensorflow implementation of LSTM and limits the number of previous seen observations. We followed Tensorflow's guidelines and biased it until the error converges during the time period contained in the hidden size of the RNN (in Figure 3 after 7 or 8 timepoints where the length of the LSTM was set to 15 previous timepoints). We instead chose to test the effect of the hidden size. The depth of the network was kept to three layers, as the spiral dataset is small, and is varied for the neural dataset. The hidden size represents the feedback arm of the RNN, and larger hidden size allows for more complex relationship with previous data observations. We show in Figure 3 bottom right, that larger the hidden size (such as 80) the loss function tends to converge in fewer training epochs. Moreover, after they are trained, these networks are more accurate with fewer datapoints (Figure 3, left), and are more sample efficient in extracting information from previous data. There are also observable differences in their accuracy, as illustrated with a sample spiral predicted after each network has observed 10 datapoints. However, the difference in prediction between 20, 40 and 80 sized networks is less pronounced especially after observing many datapoints, suggesting that regardless after many data observations the system converges roughly to the same error. This points to a certain robustness of convergence of the loss function with regard to parameter variations for long time sequences.

**Figure** Error! No text of specified style in document. **Effect of Network Size on Initial Condition Predictions**

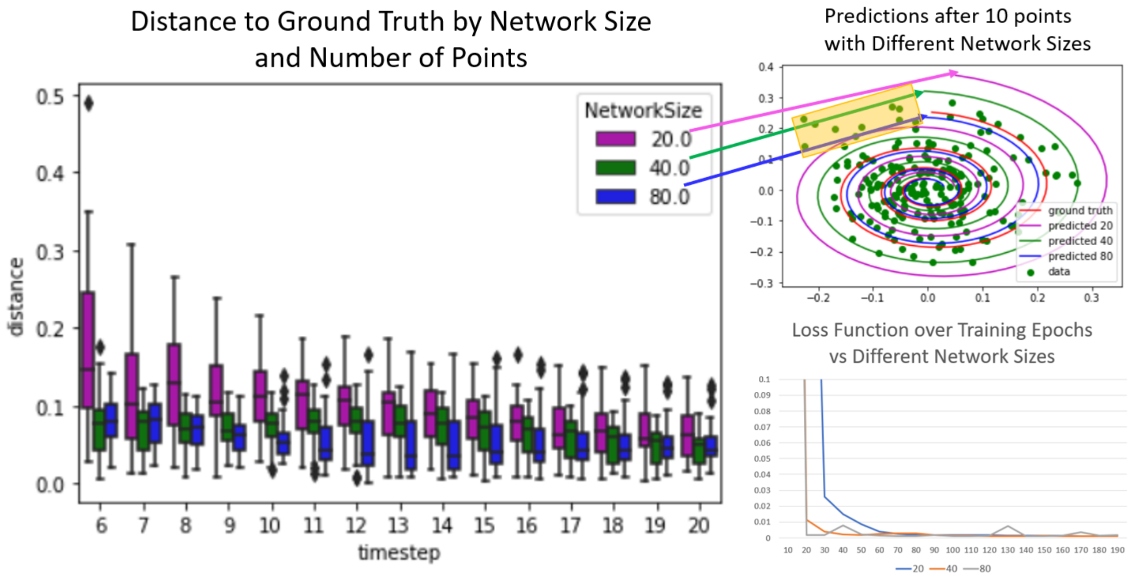


**The effect of the network size in its ability to converge to the true initial conditions with fewer samples. Three network sizes of the hidden state (20, 40, and 80) are compared while keeping the depth of the network and the number of batches constant. The plot on the bottom right shows that in all three architectures the loss function approaches zero and plateaus at roughly the same value. However, the larger networks are more sample efficient, meaning that with fewer samples, they are able to estimate better initial conditions. This is shown on the left, where the largest network (80) has the smallest distance to the ground truth. After the LSTM has viewed enough samples the differences between the networks vanishes. An example on a single spiral is shown (top right), where the predictions after seeing 10 datapoints are 0.11, 0.08, and 0.04 apart from the ground truth for the three differently sized networks at 20, 40, and 80 respectively. The actual pink and green data do not cross any of**

**the initial data, because the algorithm only estimates the initial conditions which are not precise for smaller networks, and since we are integrating from the initial conditions, the resulting trajectories are far away from the future datapoints which have not yet been seen by the Machine Learning Network.**

4. *RNN Parameter Estimations on fMRI data*

For the neural dataset we have a much larger state space having at least 66 state variables. Therefore, we tested a number of different parameters as listed below in the Table1. For a given BNM, the r-square accuracy was calculated for different parameters of the machine learning network. The base parameters were chosen from Kashyap et. al 2020 with 600 iterations, 50 for length of RNN, 150 for hidden size, and 4 layers and across each row in the table we change one of these parameters at a time, to see how it effects the performance of the algorithm. The accuracy doesn't change a lot when the parameters are varied as shown in Table 1 suggesting that the network is maximizing the information transferred from previous states like in the spiral example.

**Table 1 R-squared Accuracy at the 3rd timestep across different RNN Parameters**

| **Parameter Evaluated** | | | | |
|---|---|---|---|---|
| **Number of Iterations** | 200 | 600 | 800 | 1000 |
| | 0.729 (0.02) | 0.743 (0.021) | 0.739 (0.022) | 0.744(0.02) |
| **Length of RNN** | 30 | 40 | 50 | 60 |
| | 0.741(0.03) | 0.744 (0.028) | 0.743 (0.021) | 0.741(0.024) |
| **Hidden Size** | 100 | 120 | 150 | 180 |
| | 0.733(0.02) | 0.734 (0.02) | 0.743 (0.021) | 0.741(0.022) |
| **Number of Layers** | 3 | 4 | 5 | 6 |
| | 0.738(0.02) | 0.743 (0.021) | 0.739(0.02) | 0.730(0.02) |

**Evaluation of network parameters of the RNN. The mean r-squared (standard deviation) are given for different sized networks on the Firing Rate Model (k=0.6) on the third timestep prediction for 2500 short resting state fMRI trajectories.**

5. *Session Convergence and Metric Choices on fMRI data*

The average loss over 10 epochs each with 50 timepoints is plotted in Figure 4 left. The epochs are contiguous over time, where the hidden state of the first epoch is the input for the second epoch. The loss is the mean squared error between the predicted and the next timepoint. The first batch has a large error when the RNN is not properly initialized but then converges to a minimum, similar to the spiral dataset after observing enough timepoints. The first epoch is ignored, and all the calculations are made after the first 50 timepoints.

Once trained, instead of using the mean squared error as used in the training, we use the more general r-squared metric in order to test how well the trajectories originating at the predicted initial condition fit the future datapoints. Unlike the mean squared error metric, the r-squared metric would generalize even when the number of brain regions are changed or under different normalizations allowing it to be more compatible with future algorithms that test short term predictability. In Figure 4 right, we also tested how well the measure generalizes from testing once every epoch (50 timepoints) to testing every timepoint as well as the effects of computing the r-squared over a batch of data consisting of 60 individuals vs testing each individual at a time. The accuracy at the 3rd timepoint from the initial condition is plotted for these four conditions (i.e the permutations of individual vs group and one timepoint vs all timepoints). There is no difference in the mean or the variance in testing at every timepoint vs testing on all timepoints. This is not surprising as the algorithm was developed to predict the correct the initial conditions at every timepoint of the timeseries and the experiment shows that it generalizes and performs relatively similarly on all timepoints. On the other hand, there is a difference in the variance but not the mean when comparing the group r-squared values vs the individual r-squared values. This can be explained as individual differences in fMRI are averaged out in the group metric. This shows that our approach might be sensitive for individual differences, but we use the group metric for the subsequent results as they are more robust and allow us to test our hypothesis on the differences between different BNM. Moreover, since we utilize only a group averaged structural matrix, we are more interested how well the BNM fit to the group than to any particular individual.

**Figure 4 Individual Variability and Generalization Across Time**

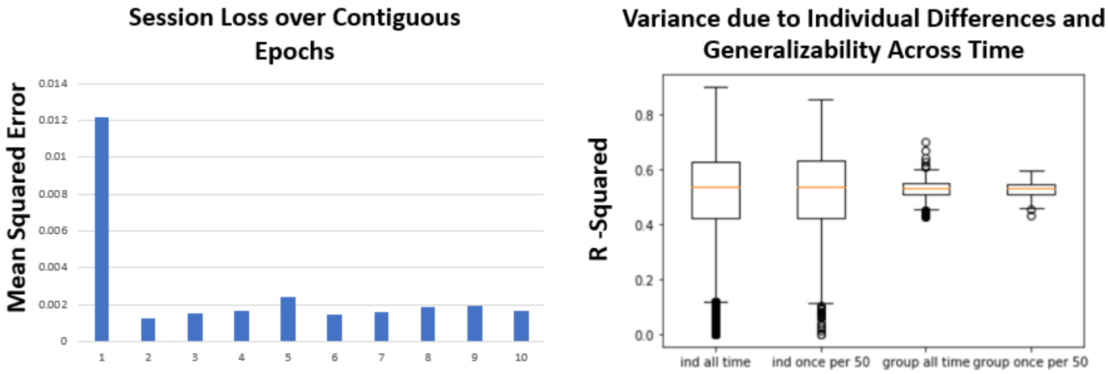


**Left: The mean squared error calculated over each epoch of 50 timepoints from a continuous 1200 timepoints of fMRI data. The loss is constant after the first epoch which is much higher since the RNN hidden state is not properly initialized. The constant loss also suggests that after the first epoch the estimation of the initial condition is constant and has converged. The first epoch is not used in any of the subsequent estimates of evaluating the dynamical system. Right: The R-squared of a FRM (k = 0.9, $\sigma$ = 0.3) is computed using 4 different methods. The first one (ind all time) evaluates the most number of tests, where the r-squared of each individual is calculated on every timepoint after the first epoch. The second one evaluates the r-squared of each individual fMRI once per epoch. The last two averages the r-squared across a batch of individuals at every timepoint and once per epoch. There is no difference in evaluating once per epoch or at every timepoint. There is a difference between the individual and group measures, which is expected as the group measure averages out the effect of individual variance. In our subsequent results we use the group measure, as our**

**model does not take into individual differences in the structural matrix, and the group measure is more robust in evaluating the differences in the ODE which is what we are interested in.**
